# Intrinsic timescales of sensory integration for motion perception

**DOI:** 10.1101/305409

**Authors:** Woochul Choi, Se-Bum Paik

## Abstract

A subject-specific process of accumulation of information may be responsible for variations in decision time following visual perceptions in humans. A detailed profile of this perceptual decision making, however, has not yet been verified. Using a coherence-varying motion discrimination task, we precisely measured the perceptual decision kernel of subjects. We observed that the kernel size (decision time) is consistent within subjects, independent of stimulus dynamics, and the observed kernel could accurately predict each subject’s performance. Interestingly, the performance of most subjects was optimized when stimulus duration was matched to their kernel size. We also found that the observed kernel size was strongly correlated with the perceptual alternation in bistable conditions. Our result suggests that the observed decision kernel reveals a subject-specific feature of sensory integration.

## Introduction

Perceptual decision making is the act of choosing an option based on the evaluation of sensory evidence ^1^. To understand how the brain translates the interpretation of sensory information into behavior, it is essential to study the mechanism by which this psychophysical judgment process occurs ^2–4^. To address this issue, human behavior in visual tasks such as motion detection has been studied extensively ^2,5,6^. In such studies, a net motion direction discrimination task has been freuqnelty implemented with a dynamic random dot display and observers’ response characteristics (i.e., reaction time, accuracy, decision confidence) were measured ^2,7–11^. Thereafter, neurophysiological studies examined the relationship between neural activity patterns and psychophysical behavior in monkeys, revealing a strong correlation between the neuronal and behavioral data ^2,5,7,12^. Similarly, computational models suggested that perceptual decision making arises through the integration of sensory information ^8,10,11^ and can be described by the diffusion-to-boundary process model ^9,13,14^.

Alternatively, it has been reported that perceptual decisions are affected not only by the sensory information, but also by other factors such as attention, task difficulty, and the feedback of the decision results ^1,15,16^. In addition, a number of studies reported substantial variation across the observers’ behavior, even in an identical stimulus condition. This inter-individual variability in perceptual behavior, often ignored or considered noise, has been recently studied more carefully using brain imaging techniques and individual variability appears to be related to local structure or connectivity of the brain ^17,18^. Further research is required, as the notion that inter-individual differences in perceptual decisions should be considered structural variations of neural circuits as opposed to mere statistical noise remains under debate.

A recent study on the perceptual decision making process during a motion perception task ^11^ suggested that subjective decision times reflects different profiles of evidence accumulated by each individual and showed that the bounded evidence accumulation model^13,14^ could predict subject behavior from their observed decision time. This suggests that inter-individual variability in perceptual decision time may be due to the synthesis of crucial information of the decision variable and the threshold in individuals, and may be of particular importance for those investigating the origin of inter-individual variability in perceptual behavior.

Given this, we hypothesized that if perceptual decisions reflect individual characteristics of each brain circuit, then the time course of sensory integration, known as the “decision kernel”, will be consistent within a subject, independent of instantaneous stimulus dynamics. We anticipate that this intrinsic decision kernel size may vary across subjects as the decision threshold varies and this may be an origin of inter-individual variability in perceptual behavior. Therefore, we suggest that wide variation in perceptual behavior originates from the intrinsic characteristics of brain circuits of individuals for sensory integration and that this should be considered as crucial information of subject-specific characteristics of perception.

To validate our hypothesis, we performed a series of psychophysics experiments using a coherence-varying motion discrimination task. We measured a decision kernel in each individual by estimating the response-triggered-average of a stimulus, while varying the motion coherence of the stimulus. We observed a very consistent profile of the decision kernel in each subject, independent of stimulus dynamics. Observed kernel size or decision time largely varied across subjects and accurately predicted the inter-individual variability in responses. Additionally, we found that the decision time-matched motion stimulus maximized the correct ratio of individual performance. Furthermore, we found that subjects’ characteristics of illusory motion perception was highly correlated with the observed intrinsic decision kernel. Therefore, our results suggest that an intrinsic, perceptual decision kernel is a critical factor to study sensory perception and that the inter-individual variability can be considered as a subject-specific trait from this decision kernel.

## Results

### Perceptual decision making during coherence-varying motion discrimination task

To characterize individual motion perception sensory integration, we designed a coherence-varying motion discrimination task. For a motion stimulus, random dots were positioned in a circular annulus and a certain portion of the dots were shifted to new rotated positions (clockwise or counter-clockwise) in the next movie frame. To generate a random pattern of motion ^10^, the portion of rotating dots (motion coherence, c) and a rotational direction (sign of c) were set to fluctuate randomly over time (see the Methods section for details). During the task, subjects were asked to report the direction of rotation as soon as they perceived a motion (Figs. 1a and b). To compare the perceptual decision characteristics under different conditions of stimulus dynamics, we varied the frequency of motion fluctuation (Fig. 1c, see Supplementary Fig. S1) from 0.15 Hz (F_1_; lowest) to 1.24 Hz (F_4_; highest).

**Fig. 1.**
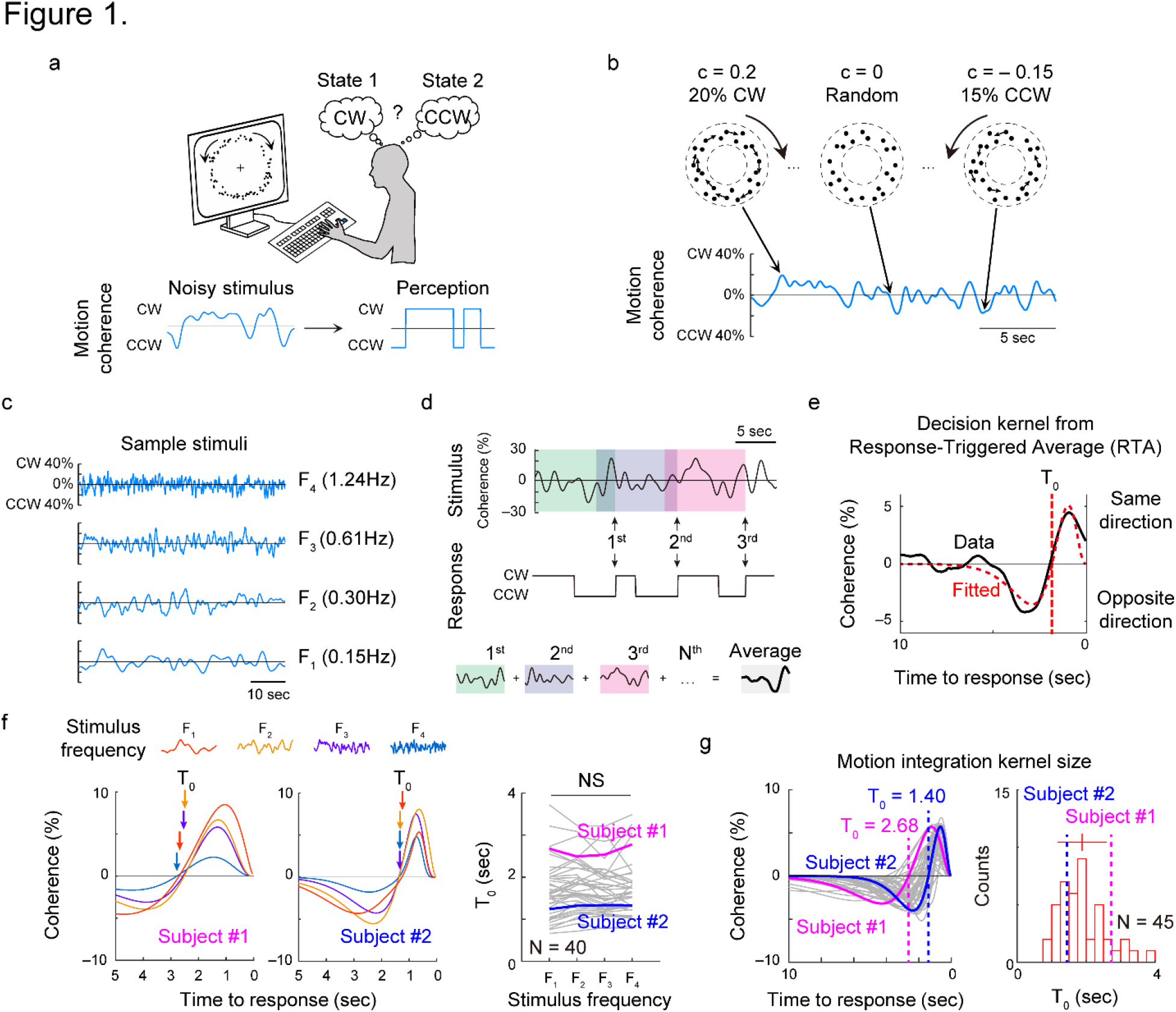
Measurement of evidence accumulation time course using coherence-varying motion discrimination task. (a) Dots positioned at random locations in a circular annulus were given as a visual stimulus. Subjects were asked to report the direction of perceived rotational motion by keyboard press. The positions of dots were updated at every 50 ms and the perceptual alternations between the two directions were recorded (b) A constant portion (motion coherence, c) of dots were controlled to rotate either clockwise or counter-clockwise. (c) Motion coherence was controlled to fluctuate with four different temporal frequencies, from 0.15Hz (F_1_) ~ 1.24Hz (F_4_). (d) At each response of motion perception (black arrows for CW switches), the preceding stimulus pattern was recorded and averaged. (e) From the observed Response-Triggered Average (RTA) kernel, the time point at which the curve becomes zero was defined as T_0_, the decision time window. (f) RTAs under four different stimulus conditions. T_0_ was fairly consistent under these conditions (One-way ANOVA, p=0.91, F(3, 156) = 0.17). (g) Fitted motion integration kernel of all subjects. Two sample RTAs were highlighted for comparison. Subject 1 (magenta) showed a longer kernel of T_0_ = 2.68 sec than subject 2 (blue) with a kernel of T_0_ = 1.40 sec. T_0_ varied from approximately 14 sec across subjects.

To quantify the subject’s perceptual decision kernel, we measured the average stimulus pattern that triggered perceptual responses using the reverse correlation method ^19–21^. We captured the stimulus pattern within the 10 second window prior to the subject reporting the direction of the perceived motion (Fig. 1d). Then, the sampled stimulus patterns were averaged together, creating the response-triggered average stimulus (RTA). The RTA measured in each subject allowed us to find the temporal profile of sensory integration for a perceptual decision, which we defined as the decision kernel of the subject (Fig. 1e). The shape of the RTA showed a positive peak before the response, which then decreased to negative value and gradually reached zero (see Supplementary Fig. S2 for control analysis). We found that an individual RTA curve fit well to a superposition of two alpha functions, similar to the quantification of the temporal receptive field structure of retinal neurons ^22^.

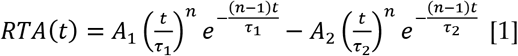

We focused on the parameter T_0_, i.e. the timing that the RTA first crosses the zero-coherence, for the profile of this decision kernel because this value reveals the size of the temporal window for effective sensory integration for decision making.

We first compared the observed RTA curves across different stimulus dynamics conditions and found that T_0_ values (the kernel sizes) were consistent across stimulus conditions, even though the frequency of motion fluctuation changed 8-fold (Fig. 1f, see Supplementary Fig. S3). We confirmed that the difference of To under different stimulus conditions was insignificant for our sample (N = 40) (p=0.91, F(3, 156) = 0.17, one-way ANOVA). This suggests that the time course of motion integration within an individual is fairly consistent and independent of the stimulus dynamics. We then averaged the RTAs from all four conditions to obtain an average motion decision kernel for each subject. In the averaged RTA, we found that the kernel size T_0_ varied noticeably from 1 to 4 sec across individuals (Fig.1g, see also Supplementary Fig. S4).

Using the observed kernels, we tried to predict the subjects’ perceptual response to the stimulus in Figure 1. From a linear convolution of the stimuli pattern and the observed decision kernel, we were able to successfully reproduce the perceptual response pattern and, in particular, N_switch_, defined as the number of perceptual switches, in each subject (Fig. 2a, see Supplementary Fig. S5). Our model predicted that the N_switch_ of the subject would be inversely related to the observed kernel size T_0_, confirmed by our observed response data (Fig. 2b and c). In addition, our model predicted that subjects with small T_0_ would have larger N_switch_ as stimulus frequency increases, while subjects with large T_0_ would have fewer changes in N_switch_ across different stimulus frequency conditions. We measured the ΔN_switch_ of each subject (Fig. 2b) and confirmed that ΔN_switch_ is inversely related to the observed kernel size T_0_, as our model predicted (Fig. 2d).

**Fig. 2.**
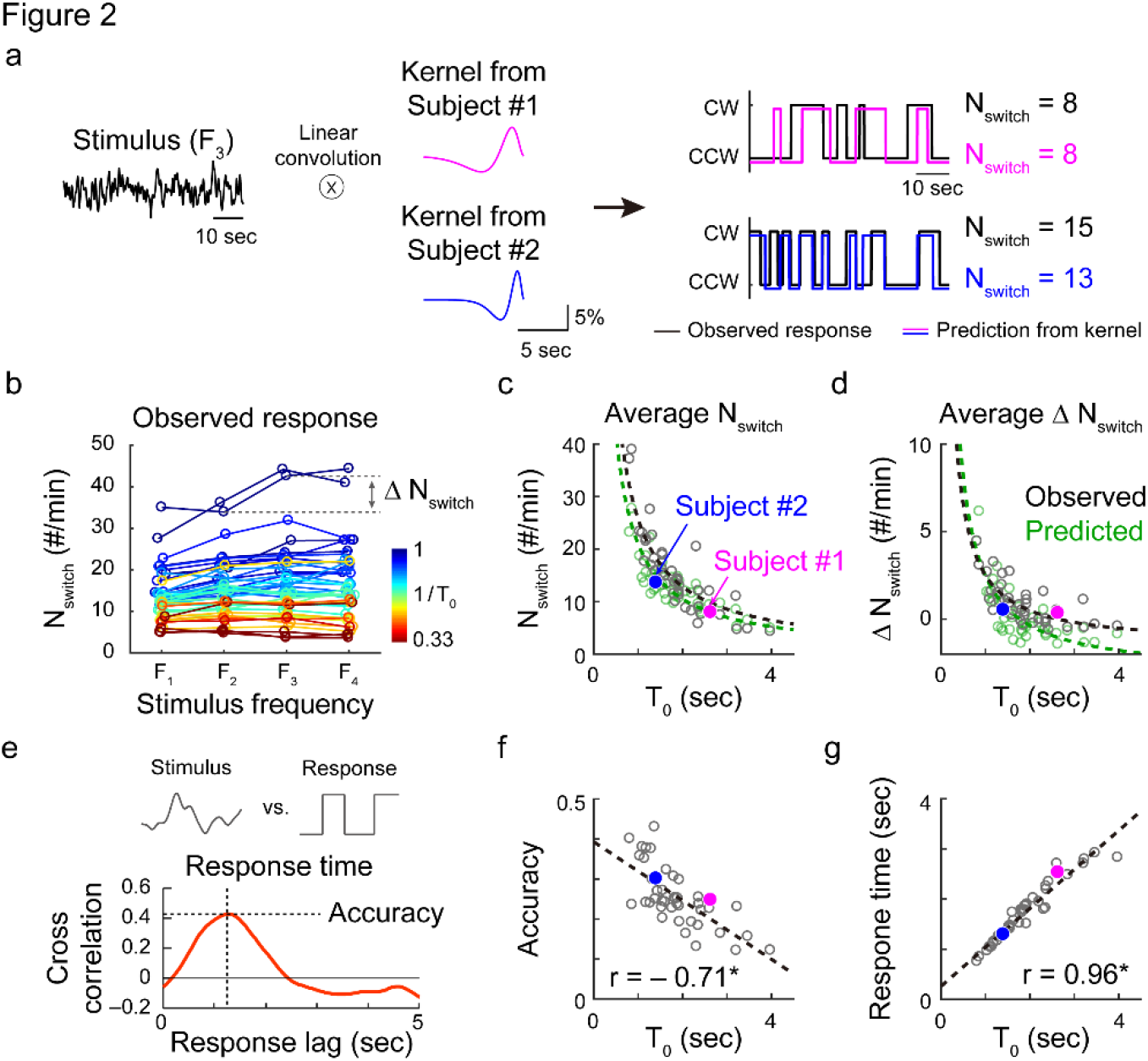
Observed motion integration kernel predicts subject’s perceptual responses. (a) Prediction of perceptual responses with observed kernels. Stimulus pattern was convoluted with the observed kernel and discretized (See Methods for details). The number of perceptual switches, N_switch_, was counted from the estimated response pattern. This prediction matched the observed responses for a given stimulus well (See Supplementary Figure S5 for details). (b) N_switch_ and ΔN_switch_ of subject responses were observed to compare with the prediction from the kernel. Each color represents data from different subjects of various T_0_. (c, d) Average N_switch_ was inversely related to T_0_ in both the model (kernel) prediction and observed data. ΔN_switch_ was also inversely related to T_0_ in the observed data, as predicted by the model. Colored filled circles show subject #1 and #2. (e) Performance accuracy and response time of subjects were defined as the maximum of cross-correlation and the corresponding time lag, respectively. (f, g) The T_0_ values of each subject were negatively correlated with the average perceptual accuracy (r = −0.71, p < 6.0×10^-8^) and positively correlated with the response time (r=0.96, p<6.7×10^-25^, Pearson’s correlation coefficient). See Supplementary Fig. S6 for details.

If the individual decision kernel size determines the number of perceptual switching during the task, we may then assume that the accuracy and the response time of each subject are also governed by the kernel size T_0_. For instance, an individual with small T_0_ may better detect the fast change of rotational direction than an individual with large T_0_. To validate this hypothesis, we defined the motion discrimination accuracy and the response time using the cross-correlation between the stimulus and response patterns (Fig. 2e). As expected, the kernel size T_0_ was negatively correlated with accuracy (Fig. 2f). Also, the response time of a subject was strongly correlated with T_0_ (Fig. 2g). These results suggest that our RTA could precisely measure the time course of perceptual decisions and the size of the temporal window T_0_ for sensory integration. We then expected that the observed subject-specific decision kernel may be responsible for inter-individual variability in perceptual behavior and might enable us to predict individual performances under a given stimulus condition.

### Kernel-matched stimulus optimizes motion discrimination performance

Based on the observations across subjects of various timescales of sensory integration, we predicted that the performance of subjects might be optimized by matching the stimulus to the observed decision kernel profile. To validate this hypothesis, we designed our next experiment to have random dots generate a motion with a fixed direction (clockwise or counter-clockwise). The motion coherence was set at a constant level (5%), but the motion duration varied from 0.5 to 5 seconds. Subjects were asked to observe the stimulus until the end of the movie and then to report the motion direction perceived at the last moment (Fig. 3a). If the accumulation of evidence is governed by the observed kernel, integrated motion information will increase as the stimulus duration increases up to T_0_, and will decrease when the stimulus duration becomes longer than T_0_ (Fig. 3b, top). Therefore, the accuracy of the perception will be the highest when the stimulus duration matches T_0_ (Fig. 3b, bottom). Our experimental results confirmed that the correct ratio did not simply increase as the stimulus duration increased, rather they showed a peak at a certain value of stimulus duration in more than half of the subjects (Fig. 3c, subjects 3 and 4). This suggests that there exists an optimal size of evidence accumulation for making the correct decision (see Supplementary Fig. S7).

**Fig. 3.**
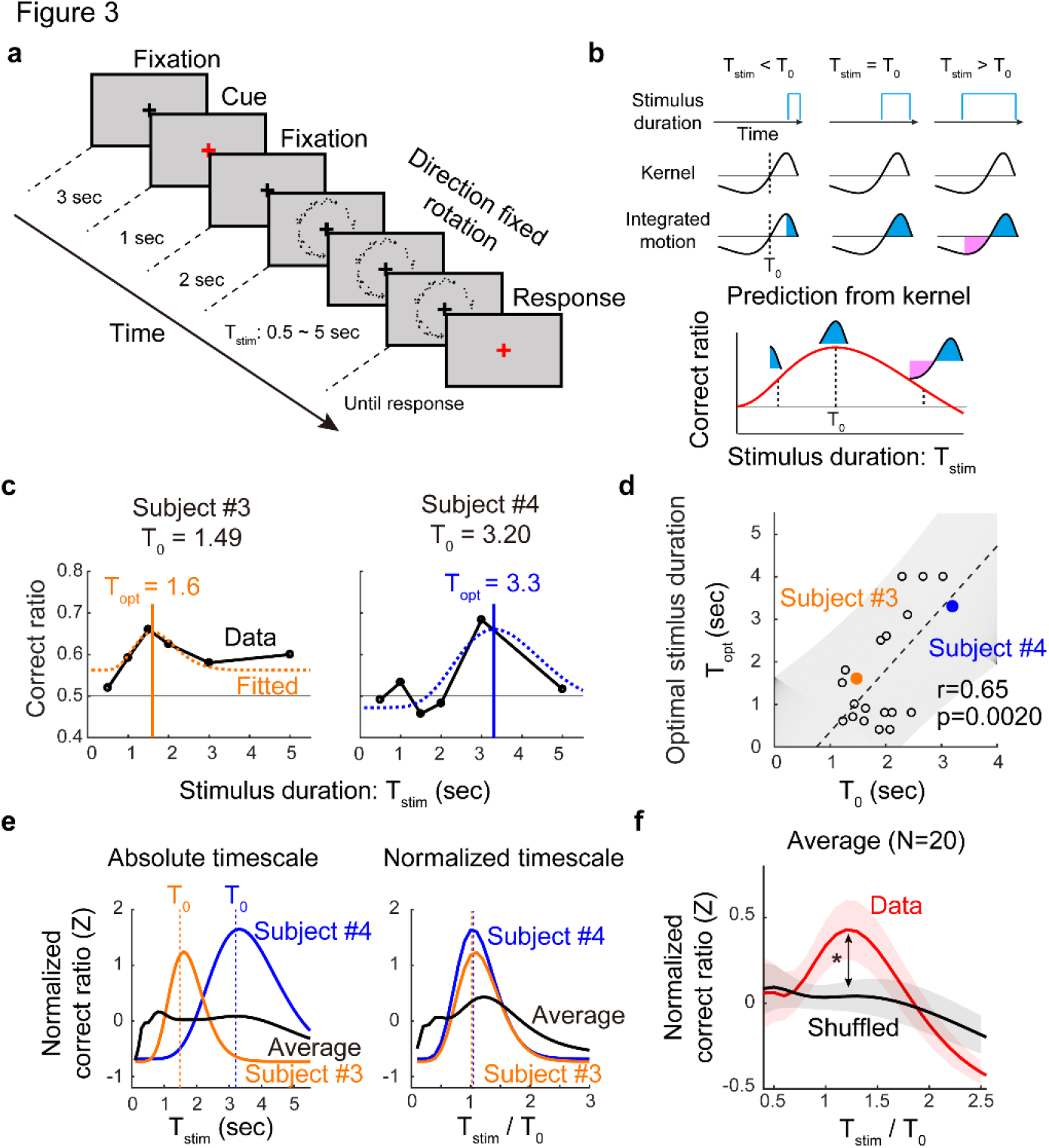
Kernel-size matched stimulus duration optimizes sensory perception. (a) Experimental design for finding an optimal value of stimulus duration. The stimulus was a constant motion of 5% coherence with fixed rotational direction and the duration was varied from 0.5 ~ 5 seconds. Subjects were instructed to report the direction of perceived motion at the end of the stimulus. (b) Correct ratio predicted from the observed kernel. Our model predicts that the integrated motion evidence would be maximized when T_stim_ matches T_0_, consequently the subject performance would show the maximum correct ratio when stimulus duration is closest to T_0_. (c) Optimal duration value at the peak correct ratio significantly varied across subjects. Two sample performance curves and their fitted value of optimal duration, T_opt_, were shown. (d) Correlation between T_opt_ and T_0_. Optimal stimulus duration was strongly correlated with the observed kernel size T_0_ (r=0.65, p=0.0020, Pearson’s correlation coefficient). Colored filled circles show subject #3 and #4. (e) In an absolute time scale, the correct ratio curves from different subjects were noticeably different (left). However, in a timescale normalized by subjects’ T_0_ value, the curves appeared to have a similar pattern with a peak near 1 (right) (f) The averaged performance curves of normalized timescale increased as stimulus duration increased toward 1 (T_stim_ = T_opt_) and then gradually decreased. The maximum correct ratio appeared at T_stim_ / T_0_ = 1.2 and was significantly higher than the control, in which T_0_ values were shuffled (black). See Supplementary Fig. S7 for details.

To examine whether the optimal perception occurs when stimulus duration is matched to the intrinsic decision kernel size, we fit the correct ratio curve to an alpha function. Then we estimated T_opt_, the stimulus duration that induces the maximum correct ratio in each subject and compared it with the individual kernel size, To. As expected, subjects’ T_opt_ was strongly correlated to To (Fig. 3d, r = 0.65, p=0.0020, N=20, Pearson’s correlation coefficient). We observed that the value of T_opt_ varied significantly across subjects, according to their decision kernel sizes. (Fig. 3e, left, orange and blue). As a result, when the stimulus duration was given as a single fixed value, each subject would show a noticeably different performance.

When we normalized the time axis of each subject’s performance curve with their intrinsic kernel size T_0_, the performance curves instead showed a similar trend, which increased toward 1 (T_stim_ = T_opt_) and gradually decreased after (Fig. 3e, right, Fig. 3f, see Supplementary Fig. S7 for details). As a result, in the normalized time scale, the population average showed a peak around 1 (Fig. 3f, red solid line), suggesting that most subjects showed the best correct ratio when the stimulus duration matched their intrinsic decision kernel size. Taken together, these results confirm that sensory integration in an individual is governed by the observed non-linear decision kernel profile and the performance of a perceptual task may also vary, depending on the difference between the kernel size and stimulus duration.

### Illusory motion perception and motion decision kernel

Thus far, our decision kernel has been estimated from apparent motion signals. We further examined the notion that the observed intrinsic kernel may predict subjects’ behavior for illusory motion perception. Previous studies have shown that random dots scattered in an annulus induce an illusory rotational motion ^23,24^ and that the perceived motion direction varies spontaneously between clockwise and counterclockwise, showing a typical bistable perception dynamic ^23,25,26^. We hypothesized that this periodic alternation in bistable perception might be also governed by the intrinsic decision kernel of subjects. To validate this hypothesis, we performed another experiment in which subjects were asked to report the direction of the perceived motion while completely random dot signals (coherence, c = 0) were shown (Fig. 4a). Consistent with previous studies, most subjects reported illusory rotational motion in this condition and the direction of perceived motion was periodically altered, spontaneously ^23^. To quantify temporal features of this bistable perception, we measured the phase duration, τ, of illusory motion in one direction. Similar to a previous report ^27^, we fit the measured τ values of a subject to a log-normal distribution and estimated the peak value 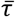, as a representation of individual dynamics of bistable perception.

**Fig. 4.**
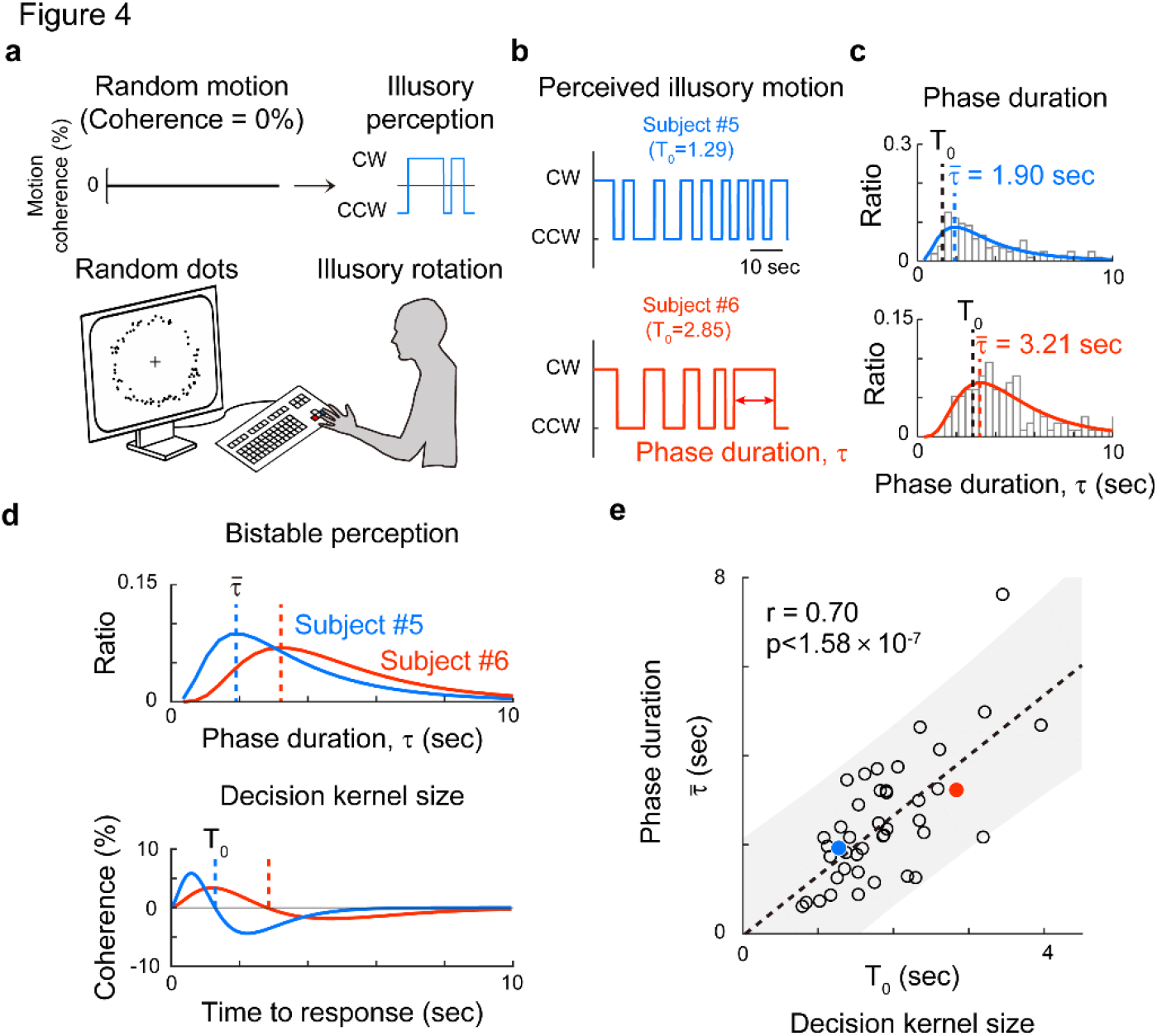
Motion integration kernel predicts the periodic alternation in bistable perception. (a) Random dot kinetics inducing illusory motion of bistable perception. Every dot is randomly distributed in each time frame, yielding no net motion. Most observers, however, perceived a rotating motion of the dots. (b, c) Sample responses from two subjects with a short (1.29 seconds, blue) and long (2.85 seconds, orange) T_0_ of integration kernel shown. In the bistable perception of illusory motion, subject 5 showed relatively faster alternation (top, blue) than subject 6 (bottom, orange) during 60 seconds of stimulation. The interval between two consecutive perceptual alternations was defined as the phase duration, τ. In each subject, the observed value of τ was fitted to a log-normal distribution and the peak value was denoted as τ. (d) The bistable phase duration τ (top) and the size of decision kernel (bottom) of subject 5 and subject 6 were shown for comparison. (e) Correlation between the τ and the size of the decision kernel. A strong positive correlation was observed (r = 0.71, p=1.58×10^-7^).

The bistable phase duration, or 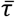, remained consistent within an individual, but varied across individuals. For example, subject 5 (Fig. 4b, top) showed relatively faster phase switching than subject 6 (Fig. 4b, bottom), but the phase durations were quite periodic and the distribution of τ values were fit well to log-normal distributions in both cases (Fig. 4c). The peak value, 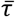, varied greatly, from 0.5 to 8 seconds across subjects (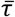 = 2.51±1.43 seconds, see Supplementary Fig. S8). However, subjects who had a long intrinsic decision time, T_0_, also tended to have slow switching dynamics with a large 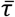, while subjects who had a short intrinsic decision time tended to have fast switching dynamics with a small 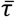. (Fig. 4d). As predicted, we observed a strong positive correlation between the values of 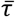 and T_0_, (Fig. 4e, r = 0.71, p = 1.58×10^-7^, Pearson correlation coefficient). This strong correlation between the observed kernel size and the switching dynamics in bistable perception suggests that the observed intrinsic decision time of sensory integration may govern the perceptual response to illusory motions, as well as apparent motions.

## Discussion

Previous studies of motion perception have suggested that perceptual decisions arise through an accumulation of evidence, thus this process can be characterized by the drift-diffusion model ^13,14^. In this bounded-evidence-accumulation model, the inter-individual variability in perceptual decisions is frequently explained by various conceptual parameters such as a decision boundary threshold, evidence accumulation rate, and choice bias ^10,11^. The model can partially predict observed experimental results such as individual accuracy of perception. However, it still remains unclear what physical variables may indeed represent those decision parameters and if any of them are intrinsically consistent to characterize individual variance of subject behavior. Our finding of an intrinsic decision kernel suggests an alternative description of the drift-diffusion model and provides direct evidence of intrinsic decision time that is subject-specific and stimulus independent. Our results also suggest that the inter-individual variability in perceptual decisions may originate from this intrinsic decision timescale and therefore may be considered a predictable trait.

We were able to demonstrate that the observed sensory integration kernel can accurately predict diverse characteristics of perceptual behavior. In our first experiment, the number of perceived motion switching under the same stimulus conditions varied across the subjects (Fig. 2b) and this number was inversely related to the observed subject’s kernel size (Fig. 2c). Moreover, it was noticeable that subjects with shorter kernel size could detect the motion direction better than the subjects with the longer kernel size when the motion coherence of the stimulus fluctuated with different frequencies (Fig. 2f, Supplementary Fig. S6). Regardless of the stimulus frequency, subjects with the shorter kernel perceived the change of motion direction better than those with the longer kernel, potentially because a shorter integration kernel may induce less sampling error in integrating noisy coherent signals than a longer sampling kernel and therefore may be advantageous for encoding highly varying stimuli (see Supplementary Fig. S6d). Another noticeable result is the strong correlation between the reaction time and the observed kernel size. In our observations, the reaction time and the kernel size were almost identical; thus the reaction time appeared very consistent within a subject and diverse across subjects, similar to the decision kernel profile (Fig. 2g and Supplementary Fig. S6). In accordance with the previous observation of the relationship between reaction time and performance accuracy, this suggests that the reaction time of a subject provides information of individual’s decision process ^11^.

Contrary to anecdotal observations, we demonstrated that longer duration of constant motion stimulus did not enhance subject performance. Indeed, when the stimulus contains a constant motion with a fixed direction, a longer duration of stimulus would generate more information accumulated in the correct direction of the decision variable, therefore the drift-diffusion model predicts a higher correct ratio of decision. In contrast, our observed decision kernel has a highly non-linear structure with a positive peak and a negative overshoot thereafter. Thus, stimulus information provided within the size of the positive part of the kernel would enhance the performance, while a longer stimulus duration may induce negative drift and degrade the decision performance (Fig. 3b). As predicted by the observed kernel, our experiments showed that there exist an optimal stimulus duration for each subject and the subject’s performance became worse when the stimulus duration became longer than this length. Therefore, our second experiment suggests that sensory integration is not a simple linear accumulation, but can be predicted by observed non-linear decision kernel within each subject T_0_ (Fig. 3e, f). This result raises an important issue; often, human psychophysics experiments are performed with fixed parameters of stimulus for all subjects and the responses are averaged across subjects to ignore inter-individual variation. Under these conditions, each subject will make a distinct decision behavior by their intrinsic kernels and the analysis could be misguided if we ignore the subject-specific traits. For example, if we simply average all the subject responses from a fixed timescale of stimuli, the averaged result may not show any clear trend (Fig. 3e, left). But, if we consider the subject-specific traits by kernel size so that the stimulus parameters were matched to the individual integration time, a common tendency of responses might be properly observed (Fig. 3e, right). This suggests that psychophysics experiments should be designed and performed carefully with a consideration of subject-specific differences.

Lastly, we showed that the observed kernel could predict the temporal features of bistable perception. The bistable perception in our third experiment is of a dynamic illusory motion, where subjects perceive a rotational motion of quasi-consistent duration from a totally random signal. For decades, it has been of interest to find the underlying mechanism of the bistable perception ^28–31^, particularly on the origin of periodic alternation of perceived states. It has been reported that the bistable switching of frequencies from different types of stimuli are correlated in each subject, suggesting a common mechanism of bistable alternation ^32–34^. Based on our results demonstrating a strong correlation between bistable switchings and the intrinsic decision time of subjects, we may argue that the observed decision kernel also governs the sensory process for the bistable condition of illusory perception. Under these assumptions, neuroimaging data in bistable perception studies may provide an insight into the origin of subject-specific dynamics of motion integration. For example, it has been reported that the structural characteristics of bilateral superior parietal lobes (SPL) were significantly correlated with the perceptual switching frequency for rotating structure-from-motion stimulus ^17,18,35^. In the functional part of the brain, both pharmacological studies and several computational models suggested that cross-inhibition levels between the two activities modulate the switching frequency of the bistable perception ^36–40^. If these factors are relevant to the observed kernel profile, it may be that individual difference of the observed kernel originate from the structural difference of the higher brain regions and the temporal scale of the decision kernel may reflect distinct inhibition level in each brain structure. Future studies should be conducted to confirm these notions.

In conclusion, we were able to verify an individual profile of sensory integration kernel from our controlled random dot stimulus and showed that human perceptual behaviors are governed by this kernel. The size of the kernel predicted an optimal stimulus duration for correct perceptual decision and the temporal characteristics of response under bistable conditions. Overall, our findings suggest that perceptual decisions arise in the intrinsic timescale of the sensory integration process.

## Methods

### Participants

Forty-five subjects (23 females, 22 males, ranging in ages from 20-29 years, with normal or corrected normal vision) were enrolled in this study. All experimental procedures were approved by the Institutional Review Board (IRB) of KAIST (KH2017-05) and all procedures were carried out in accordance with approved guidelines. Written informed consent was obtained from all subjects.

### Display and visual stimulus

Visual stimuli were presented on an LCD monitor screen (DELL U3014, 29.8 inches, 2560 × 1600, 60 Hz resolution) for all experiments. Subjects were positioned 160 cm away from the monitor and were asked to report their perception of the stimulus using buttons on the keyboard. At each frame of the stimulus, black dots were distributed in a circular annulus. The inner and outer radii of the annulus were at a 3.5 degree and 5 degree visual angle, respectively, from the center of the screen. The individual dots were 5 minute of solid angle in diameter and the dot density was set to 5 dots/deg^2^. The refresh rate of motion for each frame was 20 Hz; thus, each frame lasted for 50 ms and refreshed with the next frame. A black cross appeared at the center of the screen and each subject was asked to fix his or her eyes on the cross during the experiment. Stimulus conditions were optimized based on the results from preliminary trials and previous references ^23^. All visual stimuli were generated with MATLAB Psychtoolbox 3.0.

In the first experiment (Figs. 1, 2, and 4), subjects viewed rotating dots on the screen and were asked to report the direction of rotation by pressing the arrow keys on the keyboard whenever they perceived a change in the rotational direction of the dots (the right arrow key for clockwise rotation, the left arrow key for counter-clockwise rotation, and the down arrow key for mixed or ambiguous rotation). Subjects pushed the down arrow key for mixed/ambiguous rotation infrequently (mixed perception duration was less than 0.15% on average).

This experiment was comprised of five conditions. In one condition, the motion coherence level of the stimulus was set to 0 for a duration of 60 seconds (Fig. 4). In this condition, all of the dots in every frame were randomly located in the annulus and did not produce any global rotational motion. In the other four conditions, the motion coherence level of the stimulus, S(t), was set to fluctuate over time (Figs. 1 and 2). In these conditions, S(t) was calculated from the following equation:

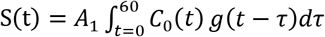

where C_0_(t) is a random number from the normal distribution of N(0, 0.05) and g(t) is a Gaussian filter:

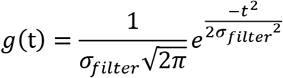

with four different σ_filter_ values of 100, 200, 400, and 800 ms. A_1_ is a constant to normalize S(t), so that the sums of absolute amplitude under the four different conditions are the same (average = 8%). The sign of S(t) determined the rotation direction (clockwise for positive, counter-clockwise for negative values). At each frame, dots of S(t) were rotated by an angle θ_rotate_ = ±5° in the next frame. The detailed statistics of S(t) are shown in supplementary Fig. S1.

In the first experiment, each subject performed a total of 80 sequences of the trials: 64 trials (16 trials×4 frequency conditions) of a coherence-varying motion condition and 16 trials of a random motion condition (S(t)=0), with random assignment of the sequence of conditions. In the second experiment (Fig. 3), the dots were set to have a fixed rotational direction, clockwise (CW) or counter-clockwise (CCW), which lasted for T_stim_. During T_stim_, the coherence level was fixed at 5%. After the visual stimulation, subjects were asked to report the rotational direction of the stimulus perceived at the last moment of the stimulus. Stimulus duration, T_stim_, was randomly chosen from the pool [0.5, 1, 1.5, 2, 3, 5] seconds (Fig. 3a). For the second experiment, each subject performed 50 perceptual decisions under 6 conditions of varying stimulus duration (300 total trials), with random assignment of the sequence of the conditions.

### Analysis

#### Motion integration kernel: Response-Triggered Average

To extract a subject’s motion integration kernel, we first measured the time point at which the perceptual switch was reported, t_switch_. In a single frequency condition, F_*i*_ of motion coherence fluctuation, we extracted the stimulus pattern 10 seconds prior to every j^th^ response of switching time, t_switch=j_ and averaged these response-triggering stimulus patterns as follows:

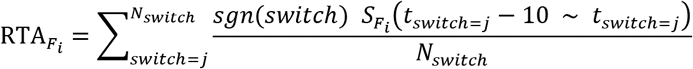

To obtain the average integration kernel of a subject, the RTAs from four different frequency conditions were summed:

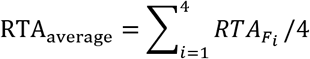

To minimize the possibility that the long and short RTAs came from the difference in switching numbers during the experiment, we generated a control response in which the responses were shuffled at random times, but with the same distribution of inter-response-interval. Then, the power of the kernel, P(t) = Σ (RTA(t)^2^) between the actual observed RTA and control RTA were compared (see Supplementary Fig. S2 for details).

#### Response prediction with observed kernel

To predict a perceptual response to a given stimulus, we took a linear convolution of the stimulus pattern with the individual motion integration kernel:

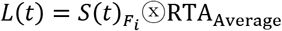

where 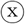 denotes the convolution and L(t) is the linear response to the stimulus.

We assumed that the response switches when the integrated response L(t) exceeds the threshold value, L_th_ were as following:

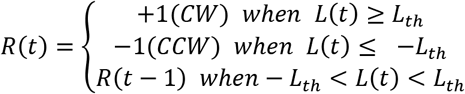

and the threshold value L_th_ was calculated from the observed kernel as:

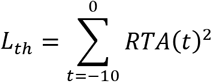

To examine the goodness-of-prediction, the cross-correlation between the R_Predicted_ (t) and the R_Observed_ (t) was calculated (see Supplementary Fig. S5). As a control, the perceptual response was switched at random times, while maintaining the same inter-response-interval of the actual response.

#### Estimation of perceptual switching of motion

During 60 seconds of a single trial, the subject’s switch responses (CW to CCW; CCW to CW) were counted (Fig. 2a) at each of the four frequency conditions. We fit the relationship between the N_switch_ and T_0_ to N_switch_ = 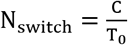, and C was estimated as 25.7 for the observed response and 20.1 for the response predicted from the estimated kernel (Fig. 2c). Also, ΔN_switch_ = N_switch;Fi+1_ – N_switch;Fi_ was calculated and fit to 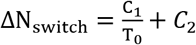 (Fig. 2d).

#### Cross-correlation between motion detection accuracy and response time

To examine the motion detection performance and response time of a subject’s behavior, the crosscorrelation between the stimulus S(t) and the response R(t) pair was calculated (Fig. 2e). Here, S(t) contains the motion coherence level at each frame and R(t) contains the simultaneously perceived state (+1 for clockwise rotation, −1 for counter-clockwise rotation, and 0 for mixed rotation). The crosscorrelation CC(t) between the S(t) and R(t) was calculated (Fig. 2e and Supplementary Fig. S6). Accuracy of the motion detection was defined as the maximum value of CC(t) at t= 0 ~ 5 seconds and response time was defined as the time lag at which CC(t) reaches a maximum value (see Fig. 2e and Supplementary Fig. S6 for details).

#### Perceptual response to a motion of different duration

In the experiment with a short visual stimulation (Fig. 3), the trial was counted as correct if the reported direction was matched the stimulus rotational direction. The correct ratio and the stimulus duration curves were fit to an alpha function:

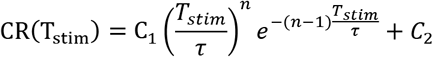

The average coefficient of determination, R^2^, was 0.5885 (see examples in Fig. 3c, and in Supplementary Fig. S7a).

In each curve of fitted correct ratio, the stimulus duration was estimated when the correct ratio reached maximum, T_opt_ (Fig. 3c). The correlation between T_opt_ and kernel size T_0_ was calculated to determine if motion integration is governed by the observed kernel. Next, we investigated the general trend of each subject’s behavior to determine whether the average correct ratio was maximized at T_0_ (see Supplementary Fig. S7). From the fitted correct ratio curve, we Z-scored the correct ratio and then rescaled the T_stim_ with respect to the subject’s kernel size, T_0_. After we obtained the normalized correct ratio curve, we averaged all subject curves. As a control, we rescaled each subject curve with shuffled T_0_ of each subject. See Fig. 3e, f, and Supplementary Fig. S7 for details.

Twenty four subjects participated in the experiment. The data from four subjects was discarded from the analysis, because their RTA and correct ratio distributions did not fit the population average, leaving a total N = 20.

#### Perceptual reponses to illusory motion in bistable condition

For the condition S(t) = 0 (Fig. 4), phase duration τ was defined as the time interval between each switch of the perceived state. For each 60-second trial, the initial 10 seconds of data were excluded for the adaptation stage and the lower 1% and upper 5% of τ data points were excluded. Measured phase durations were converted into a cumulative density function, then fit to a log-normal distribution as:

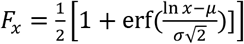

where

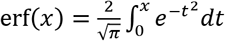

The log-normal distribution is a logarithm form of the normal distribution; thus, the peak of the τ distribution is analogous to the mean of the normal distribution. Therefore, 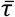 was used as the representative figure of perceptual switching distribution and 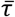 was then estimated from the fitted function as:

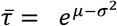

Fitting was performed using the MATLAB function ‘NonlinearLeastSquares’.

#### Statistical test

P-values and the type of statistical test used in the analysis are denoted in each figure caption and in the main text. We used a one-way ANOVA with Bonferroni correction to examine individual differences across the frequency conditions. Pearson’s correlation was used for the analysis of all linear correlations. We used a random shuffling method for comparison between the control and observed data, as described in the main text and figure legends.

## Supplementary information

Supplementary figures and legends are available in **Supplementary Information**.

## Acknowledgements

This research was supported by the Basic Science Research Program through the National Research Foundation of Korea (NRF), funded by the Ministry of Science, ICT & Future Planning (NRF-2016R1C1B2016039, NRF-2016R1E1A2A01939949) (to S.P.).

## Author contributions

W.C. designed and performed the psychophysics experiments, developed software for analysis, analyzed data, and wrote the manuscript. S.P. conceived and designed the project, directed the experiments and analysis, and wrote the manuscript.

## Competing interest declaration

Authors declare no competing interests.

---

**Fig. S1.**
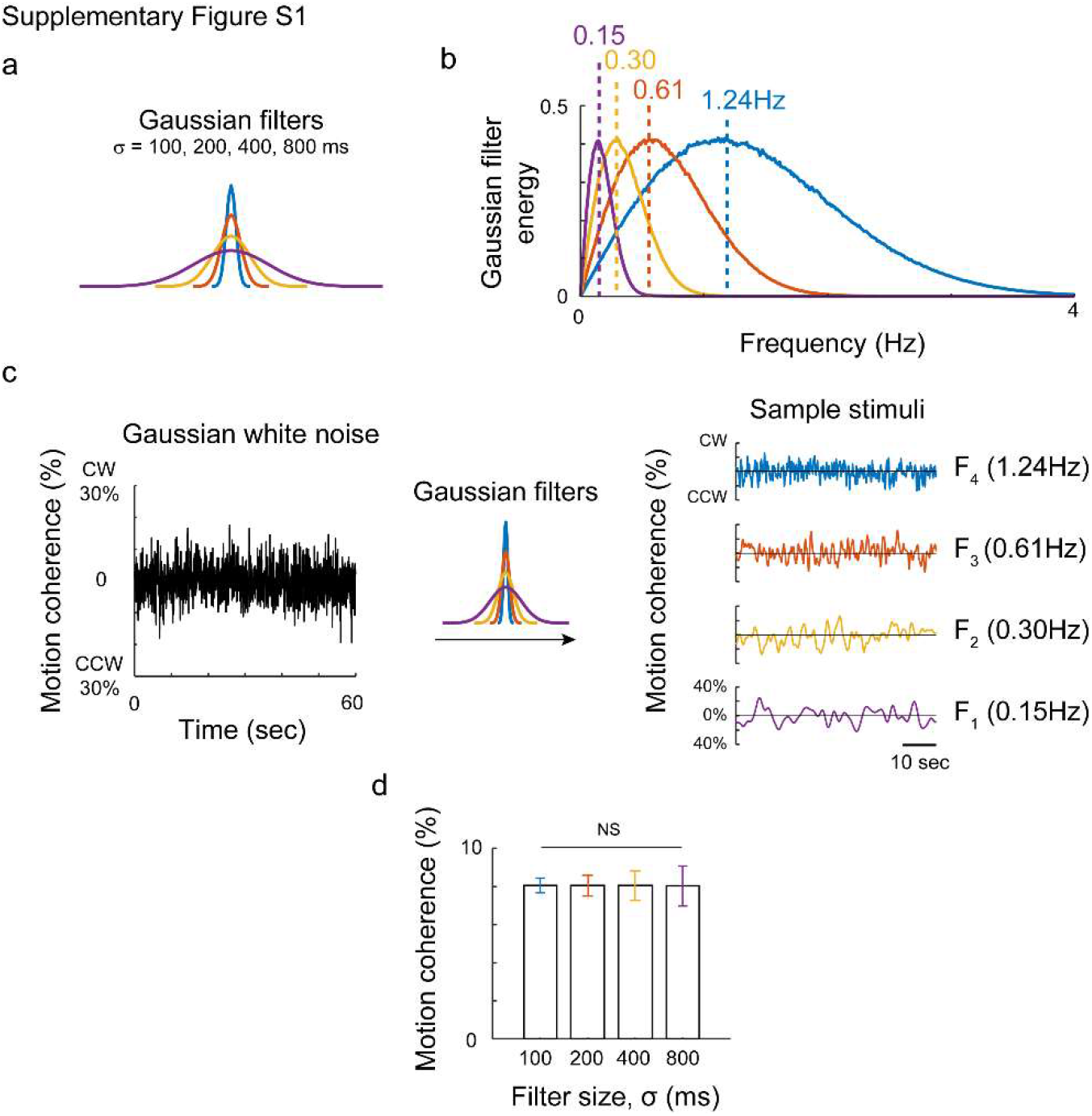
Statistics of fluctuating motion pattern. (a) Preparation of visual motion stimulus. Four Gaussian filters were used to create a time-varying motion coherence of four different frequencies. (b) The energy of the Gaussian filters in frequency space. Each filter demonstrated a peak in the frequency-energy curve, which denotes the frequency for the highest energy. The peak appeared at 0.15, 0.30, 0.61, and 1.24 Hz when the stimulus was filtered with 800, 400, 200, and 100 ms Gaussian filters, respectively. (c) Gaussian white noise was generated in every frame (left) and convoluted with a Gaussian filter with different width. (d) In these four conditions, the average coherence was normalized to have the same value (8%, N=1000 simulations, one-way ANOVA, p=0.91). Note that the average motion strength was equivalent in all conditions, thus the four conditions had, on average, the same task difficulty.

**Fig. S2.**
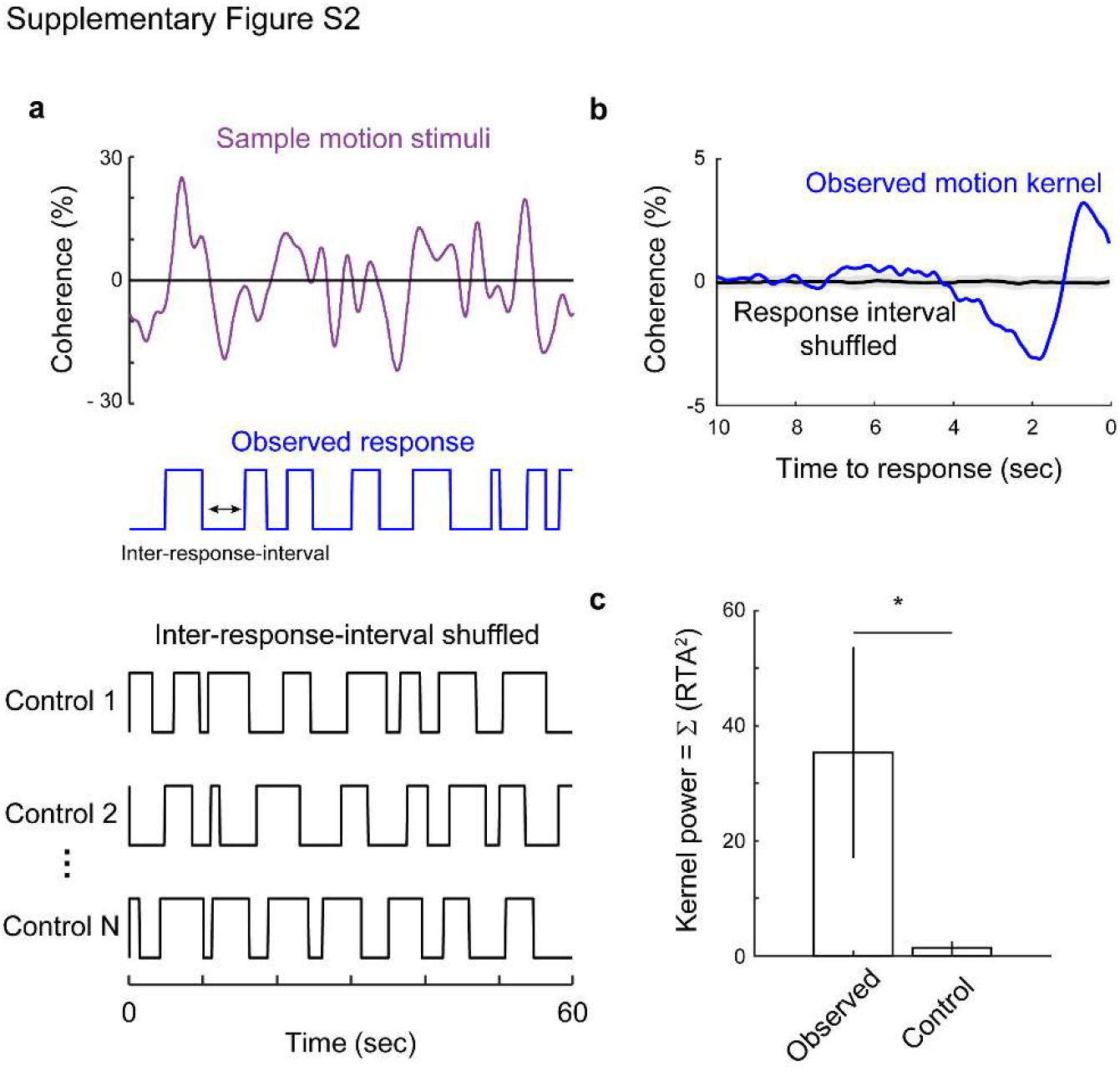
Observed motion integration kernel and a control analysis. To reject the hypothesis that the observed kernel originated from the stimulus characteristics or from the individual variance of frequent/sporadic responses, we designed a control analysis. (a) With the stimulus used in the experiment (top, purple) and the observed response (middle, blue), we made a shuffled response maintaining the same inter-response-interval of the response (bottom, black). (b) We extracted the RTA from the observed response (blue) and control response (black). The observed kernel showed a significant peak in the curve, while no peaks were found in the control RTA kernel. Shaded area denotes the standard deviation of control RTA. (c) The control RTA from the same number of responses did not show a meaningful structure. The kernel power, defined as the sum of the squared RTA, was significantly higher in the observed RTA (p < 1.49×10^-15^, paired t-test, N= 43) than in the control.

**Fig. S3.**
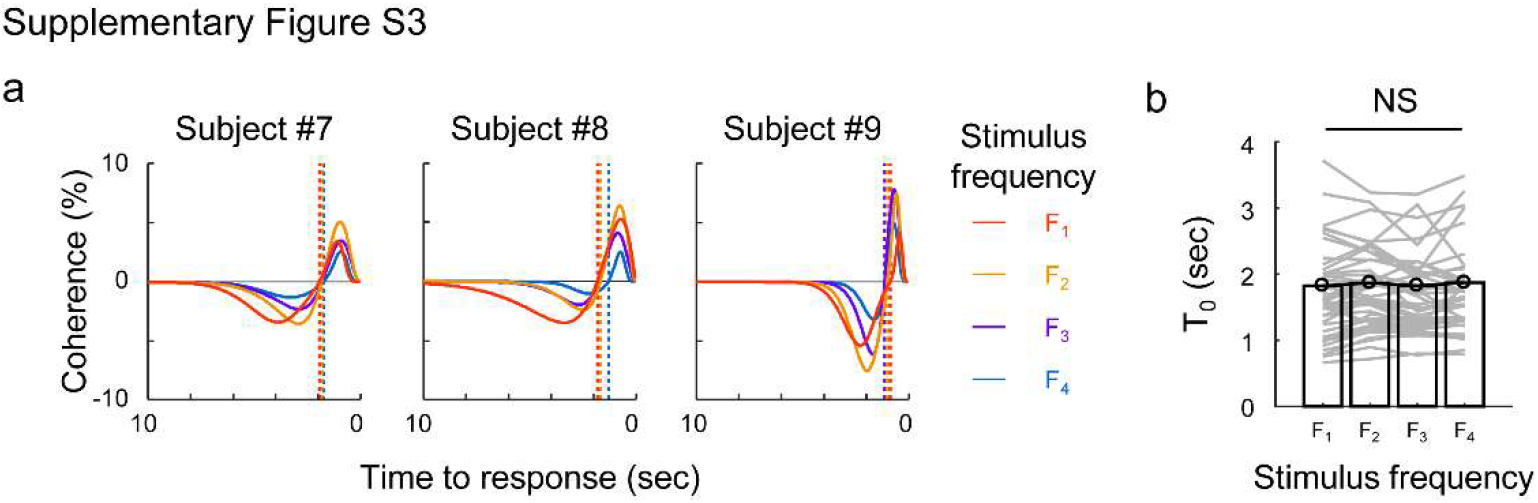
Integration kernels under four different frequency conditions. (a) Sample kernels observed from three subjects under four different conditions of stimulus frequency. T_0_, the zero-crossing point of the fitted kernel under four conditions are shown in dashed lines. (b) As shown in the Fig. 1f, a one-way ANOVA demonstrated that the T_0_ values were not significantly different in the four stimulus conditions (F(3,156) = 0.17, p=0.9143).

**Fig. S4.**
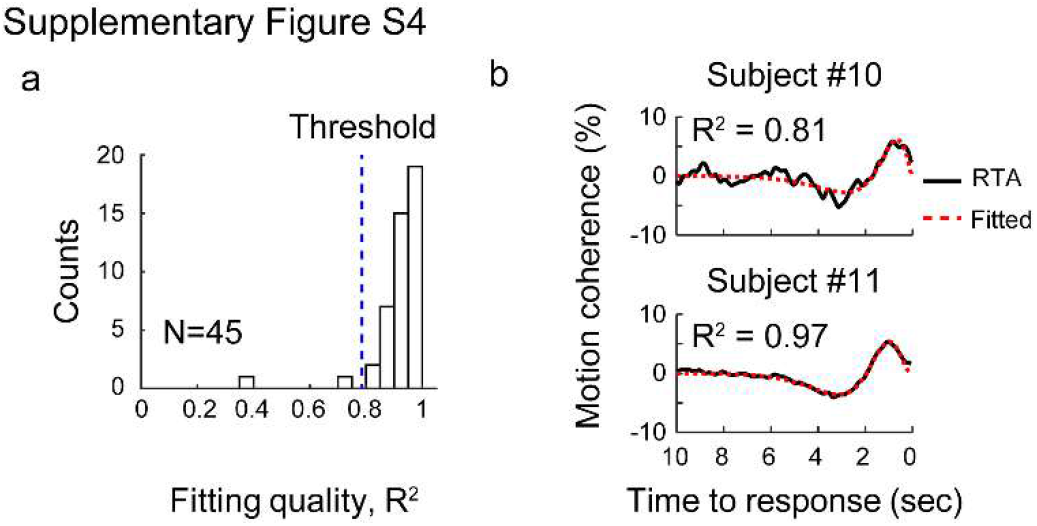
Detailed motion kernel fitting. (a) Goodness of fit of the observed kernel. The kernel was extracted for each subject (Fig. 1f) and the histogram of the coefficient of determination, R^2^, was plotted (N=45). Most subjects showed a high R^2^ (R^2^ > 0.8) but two subjects showed poor fitting result (R^2^ < 0.8), and were therefore discarded from any further analysis. (b) Sample kernel curves and fit results. The most poorly fit subject is shown in the top R^2^ > 0.8 and the most well-fit subject kernel (R^2^ = 0.97) is shown at the bottom.

**Fig. S5.**
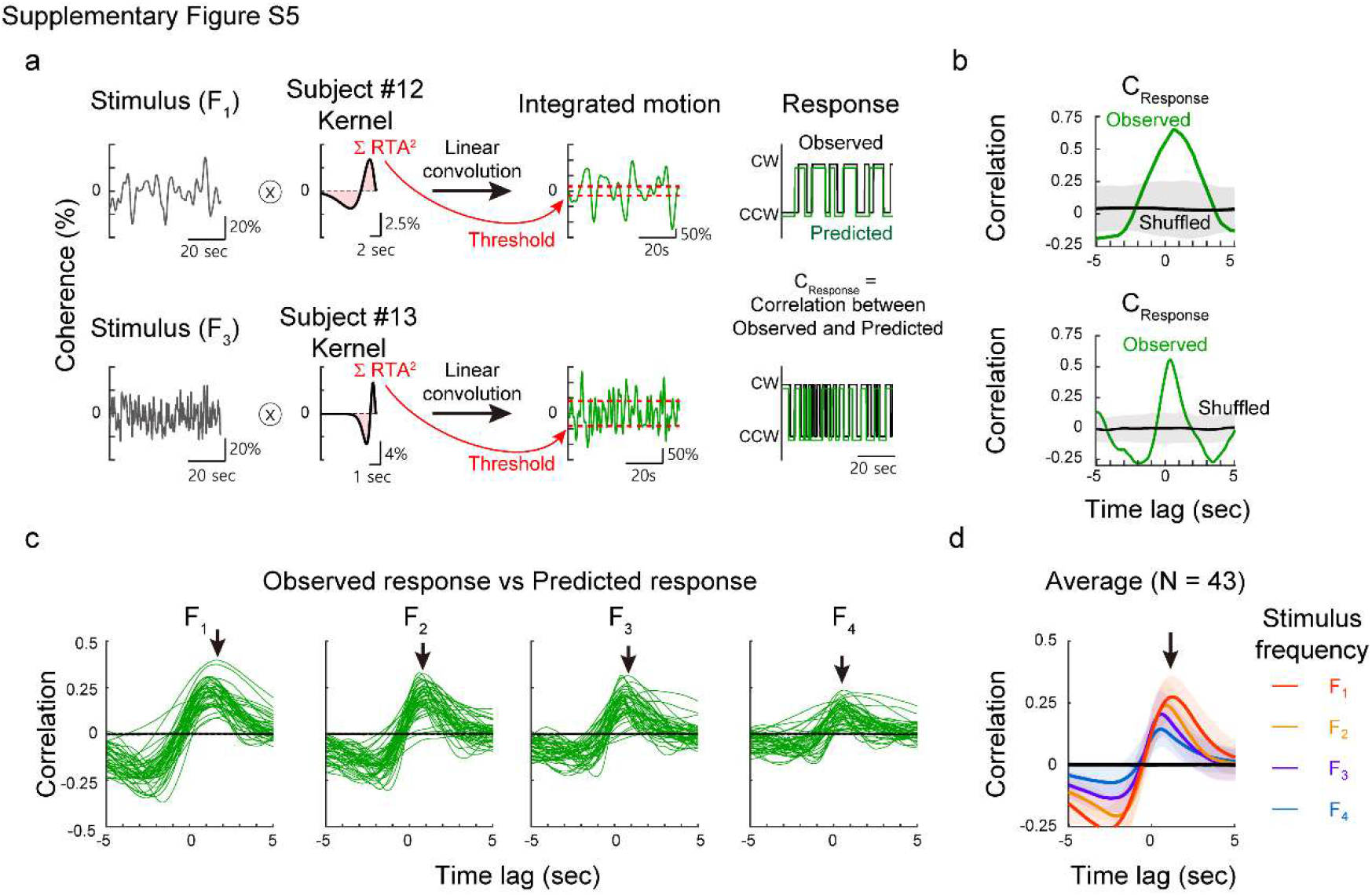
Perceptual responses predicted from the linear convolution between stimulus and observed kernel. (a) Two sample individual response predictions shown. First, the stimulus pattern used in the experiment was linearly convoluted with each subject’s average kernel (left). As a result, a predicted response curve was obtained (middle). We set a threshold value from the square sum of the kernel (red dashed line, see Methods for details), and assumed that the simulated response is switched if the linear response exceeds the threshold. We calculated a cross-correlation between the observed data (black lines) and simulated a perceptual response (green lines). (b) The model successfully replicated the observed response, which was confirmed by the high correlation value (green lines). Correlations of the time-shuffled response data was also calculated as a control (black lines). Shaded areas denotes the standard deviation of the crosscorrelation. (c) Cross-correlation of the model and observed data under four frequency conditions. Each line indicates the individual simulations. Significant peaks (black arrows) in the correlation curve showed that individual kernels can fairly well predict the response to any of the given stimuli. (d) Average crosscorrelation of the model and observed data. Each line denotes the mean correlation curve from four stimulus conditions and the shaded area shows the standard deviation.

**Fig. S6.**
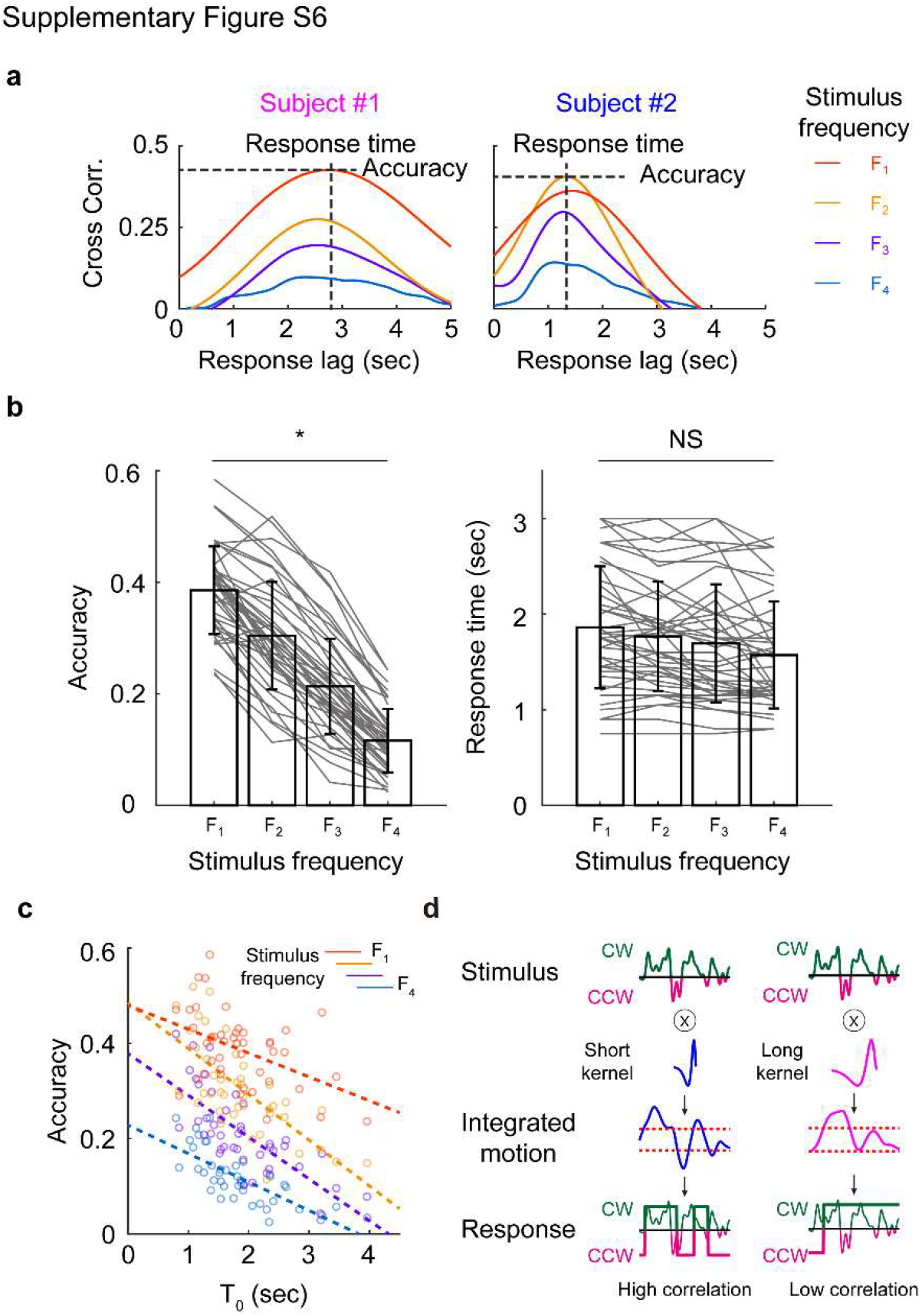
Correlation between the motion detection behavior and kernel window size. (a) Two subjects’ cross-correlation curves between the stimulus and perceptual response pattern were shown. The cross-correlation between the stimulus pattern and perceptual response was measured (Fig. 2e) under the four stimulus conditions (F1 ~ F4). The maximum amplitude of the curve revealed the accuracy of the responses; the response time was defined as the time point at which the correlation curve reaches the maximum value. (b) On average, accuracy decreased as the stimulus frequency increased (p < 1.21 ×10¯^34^, F(3, 168) = 89.49), but the response time was stable under four different stimulus conditions (p=0.15, F(3, 168) = 1.8). Estimated response time matched to the observed decision kernel size, To (Figs. 1f and 2g) (c) The accuracy of subjects under four different stimulus conditions. In all four stimulus conditions, a significant negative correlation was found between the To and the accuracy of the motion detection. (r= −0.46, −0.71, −0.74, −0.75; p < 0.0022, p < 8.70×10^-8^, p < 1.98×10^-8^, p < 5.69×10^-9^; under stimulus F1, F2, F3 and F4 conditions, respectively, Pearson’s correlation coefficient, left panel). (d) A possible mechanism for the strong correlation between the performance accuracy and T_0_. Given a stimulus (top), each subject integrates the stimulus with their intrinsic kernel. As a result, subjects with a short kernel (blue) would integrate the stimulus with a short time window and the integrated motion would change quickly (middle). Thus, the response would show a high correlation to the given stimulus. However, subjects with a long kernel integrate the stimulus with large time window (magenta), so the integrated motion would moderately follow the stimulus pattern. Thus, this subject would not follow the fast stimulus and shows a weak correlation between performance accuracy and T_0_.

**Fig. S7.**
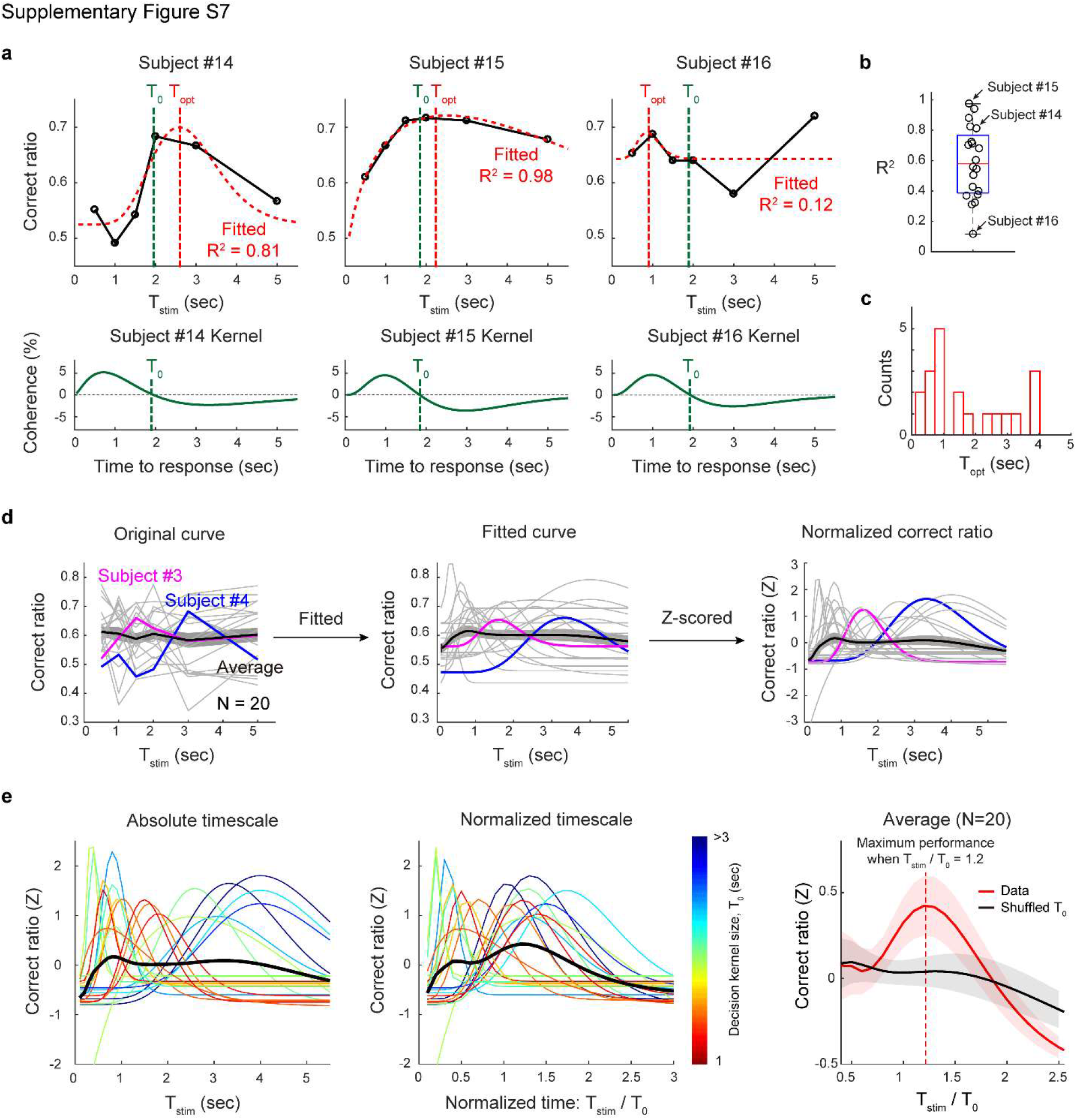
Optimized stimulation enhances the perceptual performance (a) Sample correct ratio curves and integration kernels from three subjects. Two sample correct ratio curves of good fitting subjects (14 and 15) and the curve of the bad fitting subject (16) are shown (top). T_opt_ was defined as the peak position of the curve (red dashed line), and T_0_ of each subject was shown (green dashed line). (b) The goodness of fit. The coefficient of determination is shown in the boxplot; each circle denotes the individual R^2^. (c) The distribution of T_opt_ was not biased toward the longest stimulus duration (T_stim_=5), but varied widely. (d) Normalization of the correct ratio curves. The original curve (left) was fit to an alpha function (middle) and Z-scored (right). (e) Correct ratio curve in absolute and normalized timescales. The color denotes the value of T_0_ in subjects. In a normalized time scale, the subjects had a similar trend. The population average showed maximum performance when T_stim_/T0 = 1.2 (right, red). As a control, the same correct ratio curve was normalized with shuffled T_0_ of subjects (right, black). Shaded area denotes the standard error of the mean. A paired t-test at each time point showed that the grand average was significantly different from the control at T_stim_/T0 = 1 ~ 1.6 (p < 0.05, N=20).

**Fig. S8.**
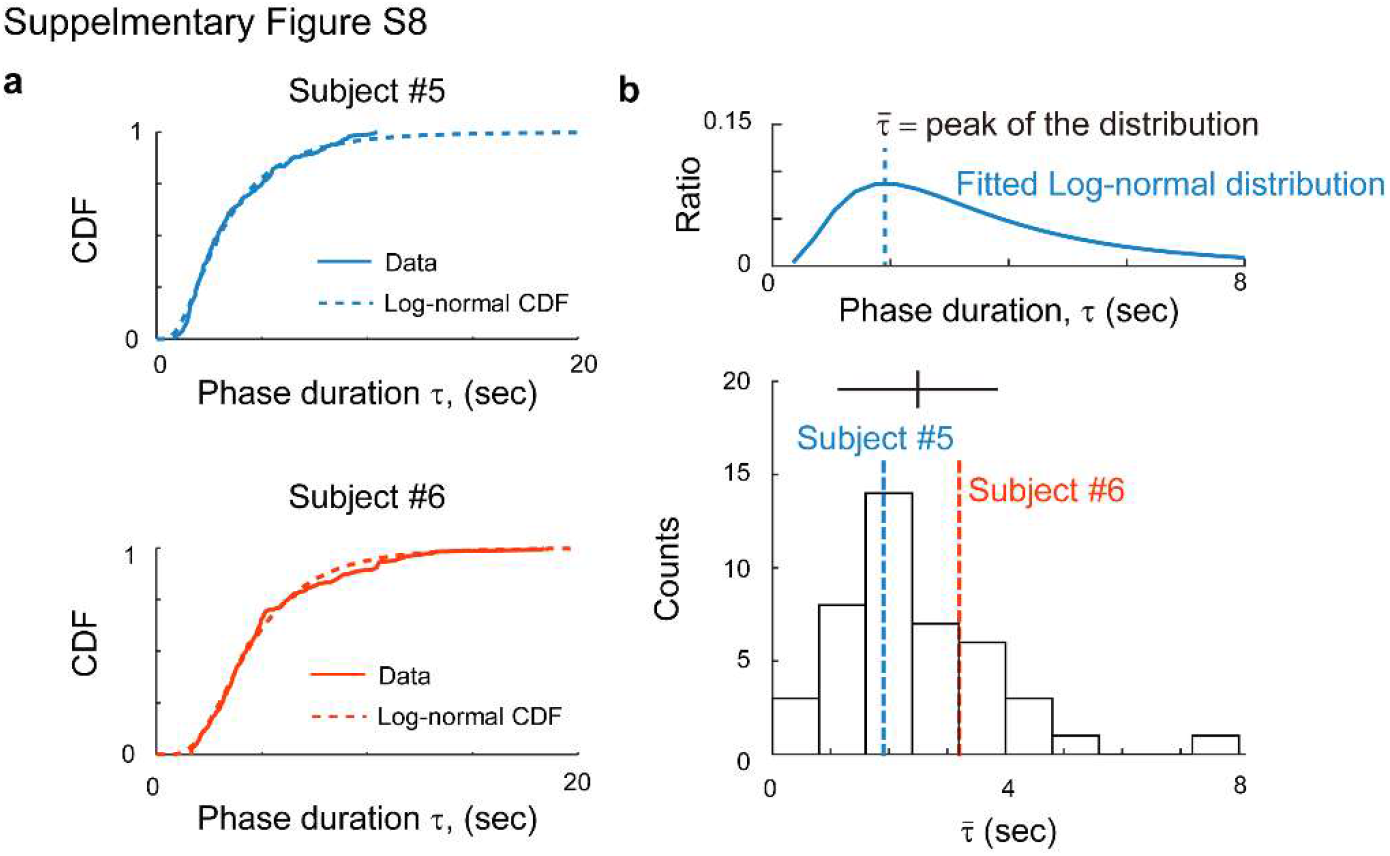
Quasi-periodic switching behavior statistics under the random bistable condition (a) The distribution of phase duration from two subjects. The τ distribution was first converted to a cumulative density function and then fit to a log-normal distribution. All subject τ distributions fit well to a log-normal distribution (Mean R^2^ = 0.92, S.D. = 0.055), demonstrating that perceptual switching occurs in a quasi-periodic manner. (b) Histogram of individual τ statistics. The peak value, 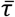 varied from 0.5 to 8 seconds, while 90% of the subject’s 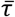 values fell between 0.69 and 4.7 seconds. The population average and the standard deviation are shown with black solid lines.

